# Regulation of mRNA transcripts, protein isoforms, glycosylation and spatial localization of ACE2 and other SARS-CoV-2-associated molecules in human airway epithelium upon viral infection and type 2 inflammation

**DOI:** 10.1101/2022.07.19.500631

**Authors:** N Stocker, U Radzikowska, P Wawrzyniak, G Tan, M Huang, M Ding, CA Akdis, M Sokolowska

## Abstract

SARS-CoV-2 infection continues to pose a significant life threat, especially in patients with comorbidities. It remains unknown, if asthma or allergen- and virus-induced airway inflammation are risk factors or can constitute some forms of protection against COVID-19. ACE2 and other SARS-CoV-2-related host proteins are limiting factors of an infection, expression of which is regulated in a more complex way than previously anticipated. Hence, we studied the expression of ACE2 mRNA and protein isoforms, together with its glycosylation and spatial localization in house dust mite (HDM)-, interleukin-13 (IL-13)- and human rhinovirus (RV)-induced inflammation in the primary human bronchial airway epithelium of healthy subjects and patients with asthma. IL-13 decreased the expression of long *ACE2* mRNA and glycosylation of full-length ACE2 protein via alteration of the N-linked glycosylation process, limiting its availability on the apical side of ciliated cells. RV infection increased short ACE2 mRNA, but it did not influence its protein expression. HDM exposure did not affect ACE2 mRNA or protein. IL-13 and RV significantly regulated mRNA, but not protein expression of TMPRSS2 and NRP1. Regulation of ACE2 and other host proteins was similar in healthy and asthmatic epithelium, underlining the lack of intrinsic differences, but rather the dependence on the inflammatory milieu in the airways.

## Introduction

Angiotensin-converting enzyme 2 (ACE2) is the major host protein used by the severe acute respiratory syndrome coronavirus-2 (SARS-CoV-2) for entering various host cells^1^. There are six *ACE2* mRNA transcript variants reported to date, which may be translated into four distinct protein isoforms^2^ **(Fig. 1a,b and Supplementary Fig. 1).** Long ACE2 isoforms 1-3 contain protein domains responsible for its physiological functions in renin-angiotensin axis and holds sites responsible for its interactions with SARS-CoV-2^3^. The ACE2 isoform 4, often referred as short or truncated ACE2, lacks the N-terminal part with the SARS-CoV-2 binding site. ACE2 protein isoforms are encoded by the distinct mRNA transcripts and they have been reported to be additionally regulated after translation by glycosylation^4^ **(Fig. 1 a,b)**. Functionally, only the short ACE2 mRNA transcript^5–7^, but not the other ones as suggested earlier^8^, is an interferon stimulated gene (ISG), and is increased upon viral infection, although it is not clear whether it is further reflected in protein expression. Since the beginning of the pandemics several groups^8, 9^, including ours^10^, have studied expression of ACE2 and other SARS-CoV-2 receptors and associated host molecules in different cells, organs and in various diseases in order to predict, to some extent, the impact of SARS-CoV-2 infection. However, these studies analyzed *ACE2* mRNA expression based on the single cell or bulk sequencing and microarray techniques, which did not distinguish between distinct *ACE2* transcripts. ACE2 and other receptors’ protein expression or their cellular localization have not been thoroughly assessed. Unfortunately, it led to at times dangerous misconceptions, such as perceiving the full-length ACE2 as being increased upon interferons or viral infections^8^ and challenging safety and validity of the ongoing interferon clinical trials. Therefore, it is crucial to cautiously revise previous findings and further elucidate the suggested and new mechanisms of its regulation at the mucosal sites.

**Figure 1.**
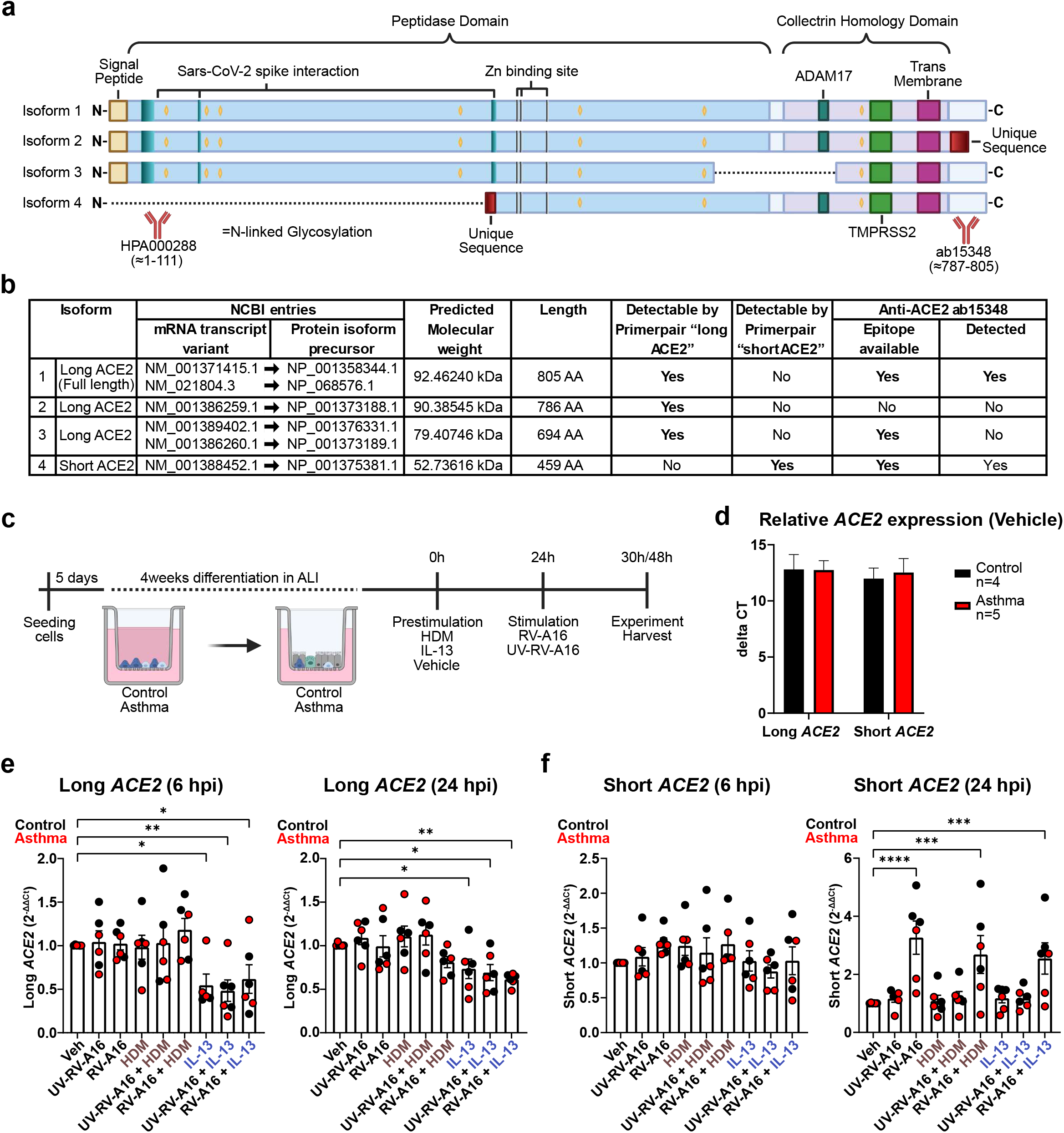
IL-13 decreases expression of long *ACE2* mRNA isoform and rhinovirus infection increases expression of short *ACE2* mRNA in primary human bronchial epithelial cells (HBEC) from healthy controls and patients with asthma. **a)** Schematic representation of ACE2 protein isoform sequences. The full-length ACE2, isoform 1, consists of an N-terminal signal peptide, a peptidase domain and the collectrin homology domain. Isoform 2 has a distinct C-terminal sequence and isoform 3 lacks the part harbouring the ADAM17 cleavage site. Isoform 4 does not possess the N-terminal signal peptide and ends in a unique sequence. N-linked glycosylation sites, SARS-CoV-2 spike interaction, Zn binding site, ADAM17 and TMPRSS2 cleavage site and the transmembrane domain are marked accordingly. Created with BioRender.com. **b)** Summary of NCBI entries of ACE2 mRNA transcripts and protein isoforms precursor IDs, predicted molecular weight, amino acid length, the ability detectability by the primers and antibodies used in the manuscript. NP_001358344.1 and NP_068576.1 lead to isoform 1, NP_001373188.1 to isoform 2, NP_001376331.1 and NP_001373189.1 to isoform 3 and NP_001375381.1 to isoform 4 respectively. **c)** Schematic representation of the experimental design. Primary human bronchial epithelial cells from healthy controls or patients with asthma were differentiated in air-liquid interface (ALI) cultures for four weeks, next they were treated with house dust mite (HDM (200 μg/ml of protein content)), IL-13 (50ng/ml), or vehicle for 24 h and later infected with RV-A16 (MOI 0.1) or treated with UV-treated RV-A16 or vehicle for 6 h and 24 h. **d)** Relative mRNA expression of long *ACE2* and short *ACE2* mRNA in vehicle-treated HBECs from n=4 controls and n=5 patients with asthma, presented as delta CT (ΔCT) with standard error of the mean (SEM). **e)** Long *ACE2* mRNA expression in HBECs from controls (black dots, n=3) and patients with asthma (red dots, n=3) upon treatment with either HDM, IL-13, or vehicle, followed by RV-A16, UV-RV-A16, or vehicle, harvested at 6 hours post-infection (hpi), left, and 24 hpi, right. RT-qPCR values were calculated by 2^-ΔΔCt^ to unstimulated condition (Veh). **f)** Short *ACE2* mRNA expression in HBECs from controls (black dots, n=3) and patients with asthma (red dots, n=3) upon treatment with either HDM, IL-13 or vehicle, followed by RV-A16, UV-RV-A16 or vehicle and harvested at 6 hpi, left, and 24 hpi, right. RT-qPCR values were calculated by 2^-ΔΔCt^ to unstimulated condition (Veh). Bars represent mean ± SEM. One way ANOVA with Dunnett’s multiple comparison correction was used to assess statistical significance. *, p < 0.05; **, p < 0.01; ***, p < 0.001; ****, p < 0.0001. **ALI**, air-liquid interface; **HDM**, house dust mite; **hpi**, hours post-infection; **Veh**, vehicle; **RV-A16**, human rhinovirus A16; **UV-RV-A16**, UV-light inactivated human rhinovirus A16.

Interactions between asthma, its viral- or allergen-induced exacerbations and coronavirus disease 2019 (COVID-19) are still unclear^11, 12^. It is suggested that allergic inflammation and the type 2 (T2)-high asthma might convey some forms of protection from SARS-CoV-2 infection, whereas T2-low asthma might be a risk factor of COVID-19^13^. It is based on the contrasting epidemiological observations in COVID-19 cohorts from different parts of the world^11, 13^, and a few mechanistic studies showing that IL-13, the major type 2 cytokine, decreases ACE2 expression and inhibits SARS-CoV-2 infection in airway epithelium^14, 15^. However, IL-13 has been also shown to be a major driver of COVID-19 severity and anti-IL-4/13 treatment is showing positive effects in COVID-19 clinical trials^16, 17^. Moreover, infection with rhinovirus (RV), the most common exacerbation-inducing factor in asthma, has been shown to inhibit SARS-CoV-2 infection via type I/III IFN-dependent mechanism^18–20^, which might suggest that this mechanism could constitute some form of protection. However, we recently reported that co-infection of rhinovirus (RV) and SARS-CoV-2 leads to a greater retinoic acid-inducible gene I (RIG-I) inflammasome-dependent damage of airway epithelium in patients with asthma^21^, which is strongly supported by the clinical findings that patients with RV and SARS-CoV-2 coinfection are more prone to severe COVID-19^22^. Finally, some components of house dust mite (HDM) or birch pollen allergens were demonstrated to decrease type I/III IFN responses^21, 23^ at the mucosal sites, a mechanism which may potentially contribute to the reported correlations between high allergen exposure and increased SARS-CoV-2 infection rates^24^.

Therefore, we aimed to analyse transcriptional, translational and posttranslational ACE2 regulation, together with its spatial localization upon allergic and viral inflammation in human bronchial airway epithelium in health and in asthma. We also analyzed other reported SARS-CoV-2 receptors and host molecules. We found that IL-13 decreased the expression of long *ACE2* mRNA and reduced glycosylation of full-length ACE2 protein via its effect on the N-linked glycosylation pathway, leading to reduction of apical expression of ACE2 on ciliated cells. RV infection increased the expression of short *ACE2* mRNA transcript, but did not change the short *ACE2* protein expression. HDM exposure did not affect ACE2 mRNA or protein. IL-13 and RV significantly regulated *TMPRSS2* and *NRP1* mRNA, but not protein. Regulation of ACE2 and other host factors was comparable in health and in asthma.

## Results and Discussion

### IL-13 decreased long *ACE2* mRNA whereas rhinovirus infection induced expression of short *ACE2* mRNA in human primary bronchial epithelial cells

In order to differentiate between the mRNA transcripts encoding for the long ACE2 (isoforms 1-3) and the short ACE2 (isoform 4), we designed primers allowing for such distinction **(Fig. 1b and Supplementary methods)**. Primary human bronchial epithelial cells (HBECs) from controls and patients with asthma were differentiated in the air-liquid interface (ALI) for four weeks. Then, we treated them with IL-13, HDM, or vehicle (medium) for 24 h, followed by the successful infection with human rhinovirus 16 (RV-A16) or treatment with UV-inactivated rhinovirus 16 (UV-RV-A16) for 6 and 24 h (**Fig. 1c; Supplementary Fig. 2a**). We found that the long and short *ACE2* mRNA transcripts were expressed in a 1^~^1 ratio at the steady-state, without any differences between controls and patients with asthma (**Fig. 1d**), therefore we analyzed them altogether in the following experiments. Long *ACE2* mRNA expression decreased upon IL-13 stimulation, whereas neither HDM exposure nor RV infection affected its expression at the measured time points (**Fig. 1e**). IL-13 did not affect the expression of short *ACE2* **(Fig. 1f).** Our findings are in agreement with the initial reports showing a decrease of *ACE2* mRNA upon IL-13^25^, with an additional notion that the IL-13 effect indeed refers to the long *ACE2* mRNA transcripts which are being translated to the functional full-length ACE2 protein, but it does not regulate the expression of the truncated *ACE2* mRNA. In contrast, we observed that short *ACE2* mRNA increased slightly already six hours post infection (hpi), which turned to notably increased levels at 24hpi (**Fig. 1f**), confirming three other reports of short *ACE2* to be an interferon inducible gene and its responsiveness to infection with the RNA virus^5–7^. Once again, our findings confirmed that the long *ACE2* transcripts encoding full-length ACE2 did not increase upon RV infection in human primary bronchial epithelium. HDM, IL-13 and UV-RV-A16 did not influence short *ACE2* expression. We did not distinguish all different long isoforms by RT-PCR, as most of the sequences share high similarity, making it challenging to find feasible primer pairs. These results underline the importance of choosing isoform specific primers for *ACE2* when reporting the results and driving relevant conclusions.

### IL-13 downregulated N-linked glycosylated ACE2, whereas rhinovirus infection did not affect the expression of any ACE2 protein isoforms

Next, we analyzed an expression of ACE2 protein isoforms in the same conditions. We used a C-terminal ACE2 antibody (ab15348), which is able to distinguish isoforms 1, 3 and 4 **(Fig. 1a)**. We detected three bands of the ACE2 protein in the human primary bronchial epithelial cells in controls and in patients with asthma (**Fig. 2a**). The band at ^~^52.5 kDa corresponded to the short isoform of ACE2 (isoform 4), while the band at 92.5 kDa to the full-length ACE2 isoform 1. We hypothesised that the ^~^130kDa band might refer to the glycosylated (N-linked glycosylation on asparagine residues as a post-translational modification) isoform 1, which we confirmed by PNGase F treatment, which led to the absence of glycosylated ACE2 (**Fig. 2b**). The absence of any band at 79.4 kDa suggests that isoform 3 is not expressed in primary HBECs. We noted that ACE2 isoform 1, glycosylated isoform 1, and short ACE2 isoform 4 were present in a 1^~^1^~^1 ratio in a similar fashion in controls and in asthma **(Fig. 2c)**. We observed a significant decrease of glycosylated ACE2 upon IL-13 treatment, whereas unglycosylated full-length ACE2 remained unchanged **(Fig. 2d, e)** with no differences between asthma and control. It suggests that IL-13 in addition to its transcriptional effect on full-length ACE2, acts also posttranscriptionally on ACE2 isoform 1. The ratio of ^~^130kDa to ^~^92.5kDa ACE2 also changed significantly in response to IL-13 **(Fig. 2f),** which suggests altered availability of unglycosylated vs glycosylated full-length ACE2. Glycosylation of ACE2 overall does not substantially influence the interaction with the receptor-binding domain (RBD) of SARS-CoV-2^26, 27^. However, a recent study showed that N-linked glycosylated ACE2 is predominantly expressed on the cell surface being accessible for SARS-CoV-2, whereas unglycosylated ACE2 is localized to the endoplasmic reticulum (ER)^28^. Therefore, our finding that IL-13 is decreasing full-length ACE2 glycosylation might translate to the lower SARS-CoV-2 infection upon IL-13 stimulation observed in other studies.

**Figure 2.**
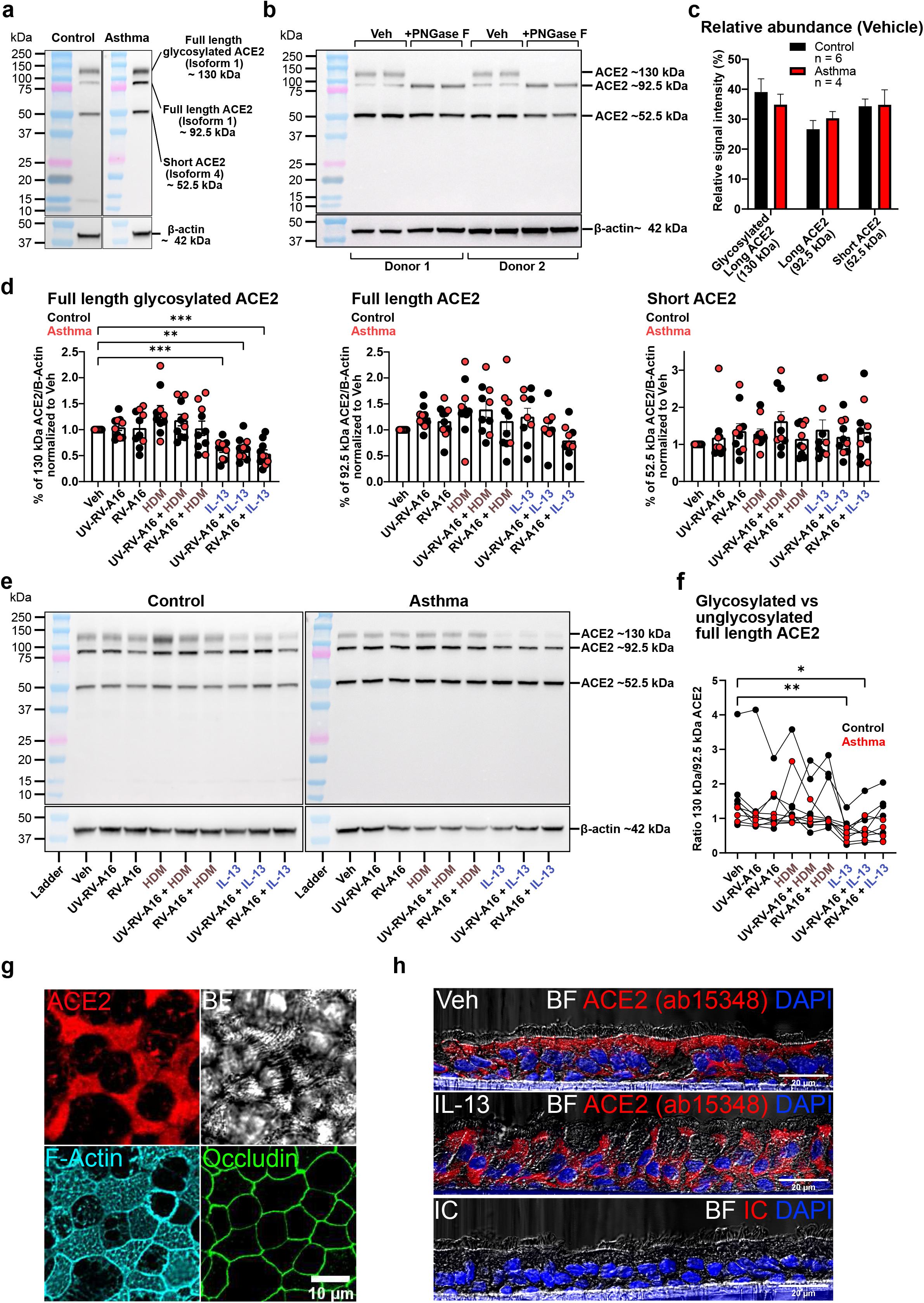
IL-13 downregulates N-linked glycosylated full length ACE2. **a)** Representative western blots of ACE2 expression in human bronchial epithelial cells (HBECs) from controls and patients with asthma. The C-terminal anti-ACE2 antibody (ab15348) revealed three distinct bands at ^~^130, ^~^92.5 and ^~^52.5 kDa, corresponding to the full-length glycosylated ACE2, full-length ACE2 and short ACE2 respectively. β-actin at ^~^42 kDa is shown in the bottom panels. **b)** Representative western blot of HBEC’s lysates from two subjects, treated with/without PNGase F revealing an absence of bands at ^~^130 kDa and more intense bands at 92.5 kDa. **c)** Comparison of ACE2 isoforms band intensities of ^~^130, ^~^92.5 and ^~^52.5 kDa vehicle-treated samples from controls and patients with asthma, shown as a relative abundance in percentage to each other. **d)** Full length glycosylated ACE2, full-length ACE2 and short ACE2 protein expression in HBECs from controls (n=6) and patients with asthma (n=4) upon treatment with either HDM, IL-13 or vehicle, followed by RV-A16, UV-RV-A16 or vehicle and harvested at 24 hpi. Intensity measurements of ACE2 were performed on western blots, standardized by β-actin intensity, and normalized to the vehicle condition (Veh). Individual values are presented in black (control) and red (asthma), the bar represents mean and whiskers SEM. One Way ANOVA with Dunnet’s multiple comparison correction was used to assess statistical significance. *, p < 0.05; **, p < 0.01; ***, p < 0.001; ****, p < 0.0001. **e)** Representative western blots of experiments quantified in FIG 2d). **f)** Ratio of glycosylated (130kDa) to unglycosylated (92.5 kDa) ACE2 in all experimental conditions. **g)** Representative confocal microscopy of apical planar view of ACE2 (detected with the ab15348 antibody) localizing to the ciliated cells in HBECs. Brightfield (BF) channel shows cilia in focus, F-actin indicates cilia in a puncta-like fashion and Occludin shows cell-cell junctions on the same field of area. Maximum intensity projection of apical Z-stacks. **h)** Representative confocal microscopy transversal view on HBECs by cryosections showing morphological alterations upon IL-13 treatment and redistribution of ACE2 signal. Maximum Intensity projection of Z-stacks. **IC**, isotype control; **BF**, bright field; **HDM**, house dust mite; **hpi**, hours post-infection; **Veh**, vehicle; **RV-A16**, human rhinovirus A16; **UV-RV-A16**, UV-light inactivated human rhinovirus A16.

The expression of short ACE2 protein isoform 4 in HBECs appeared to be very stable (**Fig. 2 e**). We did not find RV-induced upregulation of short ACE2 protein. This lack of an effect on the protein expression might suggest a high turnover of short ACE2 in bronchial epithelium, meaning that the potential increase of short ACE2 translation, mirroring an RV-induced increase of short RNA transcript, is balanced by the increase in the protein degradation. Indeed, it was shown that short ACE2 protein is very unstable and as such rarely detectable in a range of cell lines^5, 7^. Its more stable expression was demonstrated in the lung airway epithelia and liver bile duct epithelia^29^. These findings may also suggest that short *ACE2* mRNA transcript functions as a regulatory mRNA upon viral infections.

Using the same C-terminal ACE2 antibody, we found ACE2 to be localized predominantly, but not restricted to the apical side of ciliated cells (**Fig. 2g, h)**. There was also a positive signal localizing to cilia (**Supplementary Fig. 3a**), as previously reported^30^, but we found the positive signal in cilia to be rather weak by inspection of transversal sections (**Fig. 2h**). Using another commonly used N-terminal ACE2 antibody (HPA000288), we observed stronger staining in cilia (**Supplementary Fig. 3b**). However, using the same antibody for western blot, we found some clear bands at ^~^66 and ^~^54 kDa, which might suggest an unspecific binding **(Supplementary Fig. 3c)**. Transversal cryosections of bronchial epithelium revealed a substantial redistribution of ACE2 signal upon IL-13 treatment (**Fig. 2h**).

### IL-13 reduced apical ACE2, induced morphological changes in airway epithelium and altered N-linked glycosylation gene expression profile

Having demonstrated that IL-13 induces profound morphological alterations and reduced the amount of glycosylated full-length ACE2 in human primary bronchial epithelium, we searched for the underlying mechanisms of the IL-13 effect on ACE2. As we observed ACE2 to be predominantly expressed apically on ciliated cells, where it is accessible as receptor for SARS-CoV-2, we measured apical ACE2 intensities above the tight-junction layer, stained by occludin **(Fig. 3a).** Apical ACE2 intensity significantly decreased after IL-13 treatment (**Fig. 3b**), suggesting that IL-13-induced deglycosylation of full-length ACE2 led to its redistribution from the apical membrane to other cytosolic organelles^28^. IL-13 is known to induce morphological changes in airway epithelium such as goblet cells meta- and hyperplasia, reduced cilia cell numbers, as well as it leads to the epithelial barrier impairment^31, 32, 33^, all of which might play a role in reduced availability of ACE2 for SARS-CoV-2. Therefore, we also assessed the changes in apical cell shape and cell area on the planar view. In addition, the solidity and circularity of ciliated cells decreased significantly upon IL-13 treatment, while goblet cells’ area increased, and circularity decreased (**Fig. 3c**). Reduced solidity of ciliated cells in response to IL-13 might be caused by the hypertrophy of goblet cells and might also contribute to lower availability of ACE2 on the apical surface. We also observed and confirmed a more pronounced IL-13-induced opening of tight junctions in asthma, as previously reported (**Fig. 3a**)^34^.

**Figure 3.**
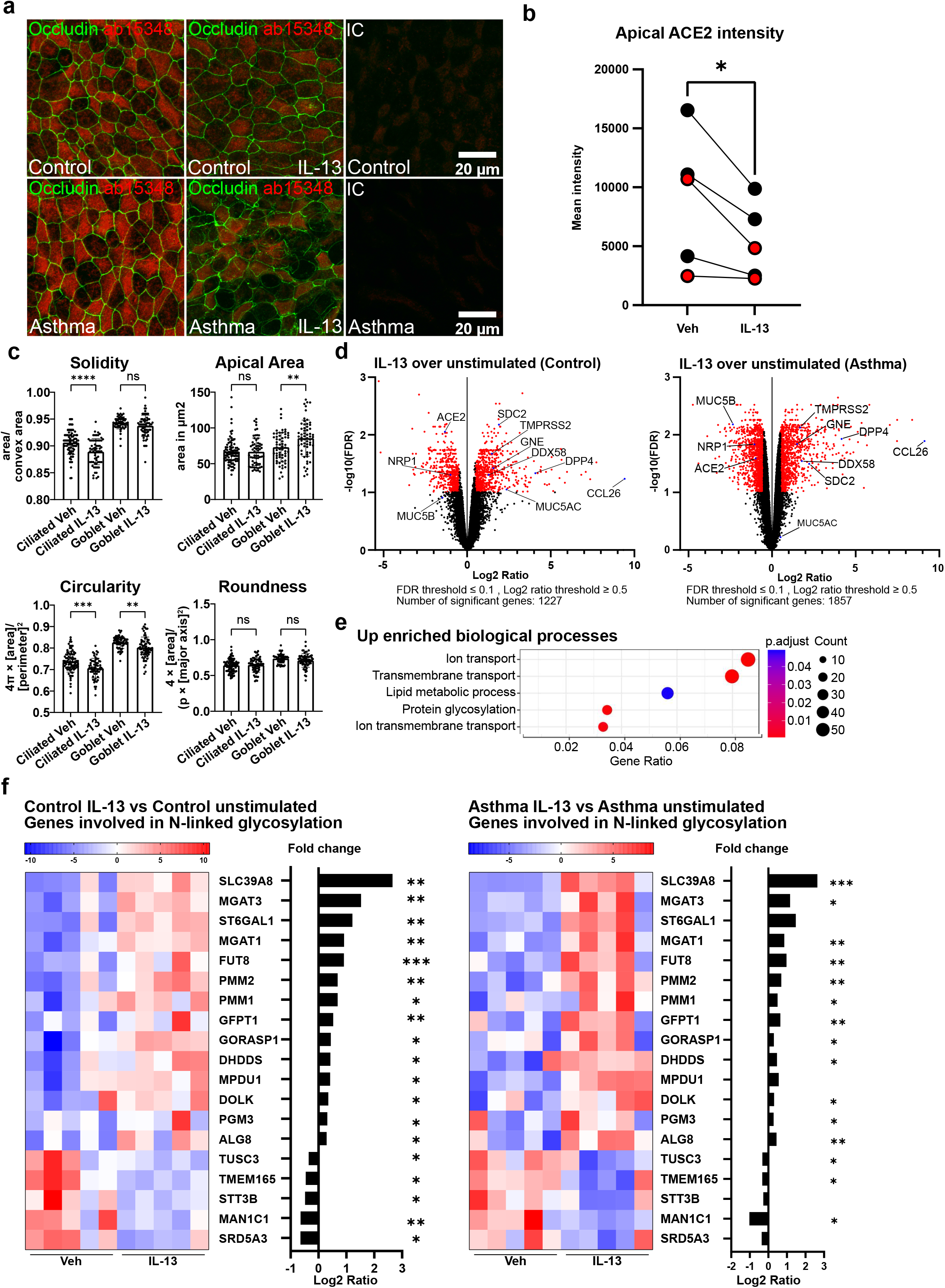
IL-13 reduces apical ACE2 by inducing morphological changes in the epithelium and altering N-linked glycosylation process. **a)** Representative confocal microscopy of apical view on HBECs from controls and patients with asthma upon IL-13 stimulation(50ng/ml, for 48h), stained with ACE2 (ab15348) and Occludin. **b)** Mean apical ACE2 intensity upon IL-13 treatment, measured on Z-projection from top up to occludin layer. Control: n=3, Asthma: n=2. Paired t-test was performed; *, p<0.05. **c)** Cell solidity, roundness, circularity and cell area measurements of ciliated and goblet cells upon IL-13 treatment. Measurements were performed on >700 cells total from 4 independent donors. Unpaired t-test was performed; *, p < 0.05, **p < 0.01, ***p < 0.001, ****p < 0.0001. **d)** Volcano plots of genes in HBECs of controls (n=5) and asthma (n=5) treated with IL-13 or vehicle, measured by bulk RNA-sequencing. Red dots are shown by a FDR value of less or equal to 0.1 and with a log2 ratio threshold greater or equal than 0.5. **e)** Top five enriched biological processes in the IL-13-upregulated genes in HBECs from control individuals. **f)** Heatmaps and fold changes (log_2_FC expression) of differentially expressed genes in N-linked glycosylation (GO: 0006487) pathway upon IL-13 treatment in controls (n=5), left and asthma (n=5), right). **IC**, isotype control; **Veh**, vehicle.

To deeper examine the mechanistic influence of IL-13 on bronchial epithelium, we performed RNA sequencing of HBECs from controls (n=5) and patients with asthma (n=5) treated with/without IL-13. IL-13 induced a significant change in the transcriptomic profile in both groups with the similar directionality of changes (**Fig 3d**). The unbiased enrichment analysis confirmed previous observations in similar models (GSE106812; GSE37693)^14^, showing that the top IL-13-affected pathways include ion and transmembrane transport, lipid metabolic processes and-, interestingly protein glycosylation **(Fig. 3e)**. Having found this, plus observing that IL-13 induced deglycosylation of full-length ACE2 and redistribution of ACE2 from the apical side of ciliated cells to the cytosol, we analyzed in more detail the N-linked glycosylation process (GO-term: 0006487). Indeed, we found that IL-13 significantly changed expression of several genes involved in this process (**Fig. 3e**), suggesting that it might be a mechanism by which IL-13 is regulating ACE2 glycosylation. Interestingly, IL-13 downregulated *STT3B* – a part of the oligosacharyltransferase (OST) complex, which was recently reported to suppress SARS-CoV-2 infection when blocked^35^. Also, *TUSC3*, which is an accessory protein of the (OST) complex was downregulated by IL-13^36^. *MAN1C1*, also downregulated by IL-13 in our dataset, has been described to be increased in severe COVID-19 patients and to be associated with the risk of disease progression^37^. We did not find any significant differences in IL-13-induced changes on N-linked glycosylation comparing asthma over control. It all suggests that IL-13 released from T helper 2 (Th2) and type 2 innate lymphoid cells (ILC2) cells at the mucosal sites during type 2 inflammation in allergic asthma on top of other known morphological and transcriptional changes might potently induce ACE2 deglycosylation and redistribution which might alter SARS-CoV-2 infection efficiency. Recent publications show that IL-13 can inhibit SARS-CoV-2 infection, as well as can reduce intracellular viral load and cell-to-cell transmission in airway epithelial cells^14, 38^. IL-13 induced mucus hyperproduction can inhibit SARS-CoV-2 infection by forming a physical barrier, but even after mucus removal, viral loads remained to be lower upon IL-13, suggesting further mechanisms to be involved^14^.

### IL-13 and rhinovirus infection regulated mRNA expression of other SARS-CoV-2-entry related molecules in bronchial epithelium, but did not change TMPRSS2 and NRP1 protein expression

Finally, as we previously analyzed mRNA expression of SARS-CoV-2-related host molecules in patients with asthma and other COVID-19 comorbidities^10^ at the steady state, here, we focused on further characterization of their expression upon viral and allergic inflammation and localization in primary bronchial epithelium. From the constantly growing list of SARS-CoV-2-entry-related host molecules (Supplementary Fig. 4c)^39^, in our RNA-seq dataset, we identified that, in addition to downregulation of *ACE2*, IL-13 also downregulated neuropilin 1 (*NRP1*) (**Fig. 4a**), which could serve as an additional SARS-CoV-2 receptor ^40, 41^. In contrast, *SDC2* (Syndecan 2) mRNA, encoding a heparan sulphate proteoglycan, was upregulated upon IL-13, which might affect the initial attachment and internalization of SARS-CoV-2^42^. Also *GNE* (Glucosamine (UDP-N-Acetyl)-2-Epimerase/N-Acetylmannosamine Kinase) was upregulated, which is crucial for sialic acid biosynthesis^43^ and potentially influences SARS-CoV-2 infection^44^. Interestingly, *DPP4* was highly upregulated in these conditions. Although DPP4 seems not to directly bind SARS-CoV-2 in humans, it continues to be controversially discussed, especially in terms of obesity and severe COVID-19^45^. Transmembrane serine protease 2 (*TMPRSS2*), a host protease facilitating fusion of SARS-CoV-2 with the cellular membranes^1^, was upregulated upon IL-13, as previously reported^25^. We and other also found *TMPRSS2* to be upregulated at baseline in bronchial biopsies of patients with asthma^10, 25^. There were no differences in the expression of these genes between controls and patients with asthma after IL-13. Functional consequences of the increase in *TMPRSS2, SDC2, GNE* and *DPP4* expression need to be further investigated, as they might play a role in the non-T2 asthma in the absence of IL-13 effects on ACE2 and NRP1.

**Figure 4.**
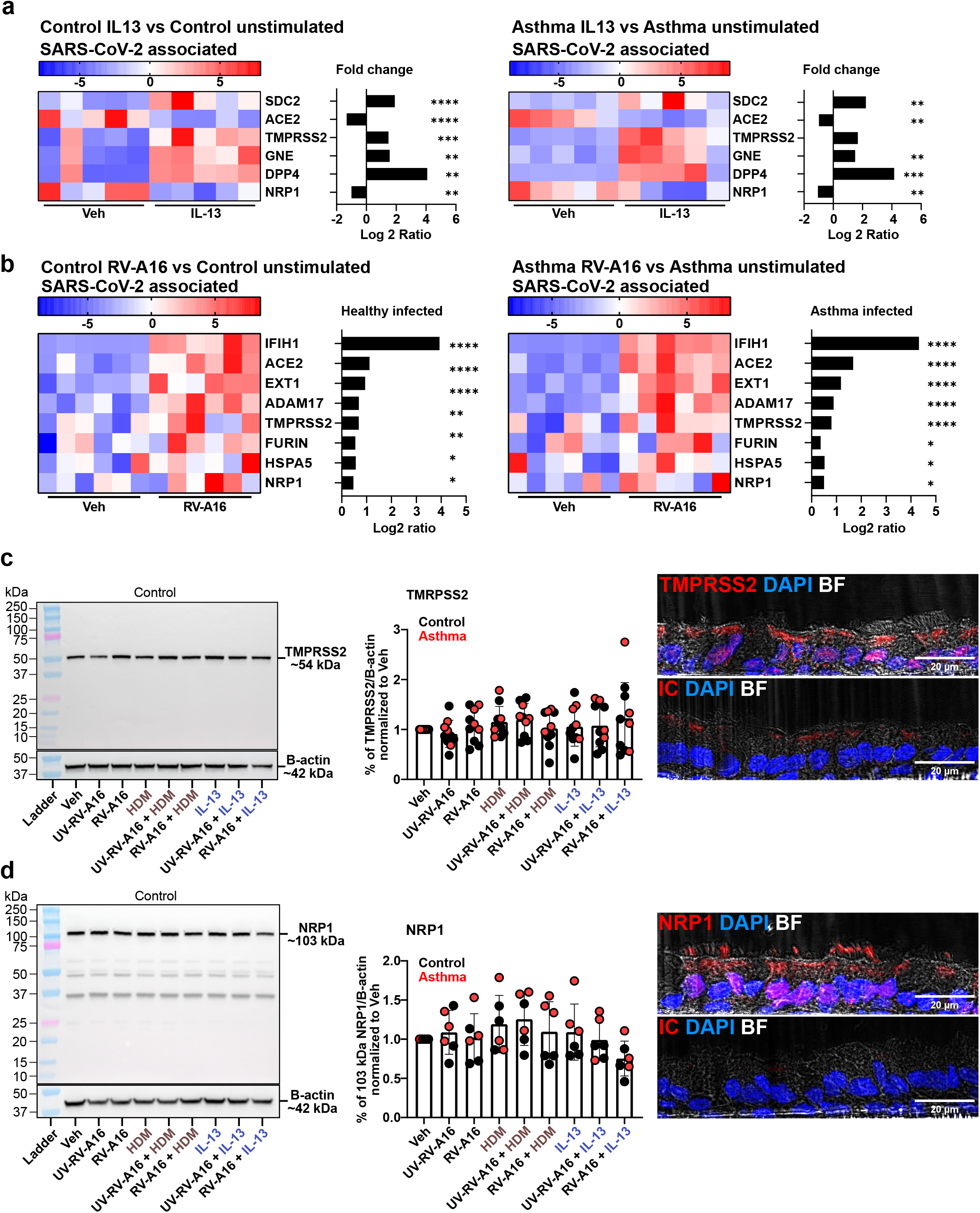
IL-13 and rhinovirus infection regulate mRNA expression of some SARS-CoV-2 entry-related molecules, but did not change TMPRSS2 and NRP1 protein expression in human bronchial epithelial cells in health and asthma. **a)** Heatmap and fold change (log_2_FC expression) of the top six significantly changed genes of SARS-CoV-2 associated molecules in controls, left, and asthma, right, upon IL-13 treatment. Control: n=5, Asthma: n=5. **b)** Heatmap and fold change (log_2_FC expression) of top significantly changed gene expression of SARS-CoV-2 associated molecules in controls (right) and asthma (left) upon RV-A16 infection. Control: n=6, Asthma: n=6. **c)** Representative western blots (left) and quantification (middle) of TMPRSS2 protein expression. Control: n=6, Asthma: n=4. Representative confocal staining of TMPRSS2 in HBECs from the control individual are shown on the right. **d)** Representative western blots (left) and quantification (right) of NRP1 protein expression. Control: n=3, Asthma: n=3. Representative confocal staining of NRP1 in HBECs from control individual is shown on the right. **c-d)** Statistical analysis were performed comparing treated samples with untreated (Veh) samples, using Dunnett’s Multiple comparison on One way ANOVA, not significant. **IC**, isotype control; **BF**, bright field; **HDM**, house dust mite; **hpi**, hours post-infection; **Veh**, vehicle; **RV-A16**, human rhinovirus A16; **UV-RV-A16**, UV-light inactivated human rhinovirus A16.

To analyze the effect of RV infection we performed data mining of previously published dataset of HBECs infected with RV-A16 (GSE61141^46^). Upon RV infection, *IFIH1* (encoding MDA5) was one of the top upregulated SARS-CoV-2-associated genes (**Fig. 4b**). MDA5 is a pattern recognition receptor which, once activated, induces an interferon response. SARS-CoV-2 RNA is recognized by MDA5^47^. Infection with RV initiates type I/III IFN response, therefore pre-infection with rhinovirus reduces SARS-CoV-2 replication^48^, but RV and SARS-CoV-2 co-infection leads to the greater damage of airway epithelium, especially in patients with asthma^21^. An observed *ACE2* increase was caused presumably by an increase in short *ACE2* mRNA, as we demonstrated above. Finally, RV infection led to an increase in mRNA expression of *EXT1, ADAM17, TMPRRS2, FURIN, HSPA3* and *NRP1*, which potentially might affect a few mechanisms involved in SARS-CoV-2 infection. All analyzed genes of SARS-CoV-2 associated molecules upon IL-13 and RV infection are shown in the **Supplementary Figure 4**.

Since *TMPRRS2* and *NRP1* mRNA were regulated by IL-13 and RV infection, we analyzed their protein localisation and expression in these conditions and upon HDM exposure. Surprisingly, we did not observe significant changes in TMPRSS2 or Neuropilin-1 protein expression in any of the experimental settings (**Fig. 4c, d**). This again suggests a high protein turnover, translation repression, mRNA degradation, or that the transcriptional change in expression might be reflected on the protein level later than 24 to 48 hours, respectively. TMPRSS2 localized apically on ciliated cells, but we could also observe positive signal localizing to nuclei (**Fig. 4c**). Similar to TMPRSS2, we detected NRP1 signal to localize apically on ciliated cells and nuclei and we found a remarkably strong positive signal in cilia **(Fig. 4d).** Altogether, we found ACE2, NRP1 and TMPRSS2 to be predominantly localized apically on ciliated bronchial epithelial cells, which makes these cell-types being the most susceptible to SARS-CoV-2 infection.

A limitation of our study is that the shed or soluble ACE2 was not investigated. In addition, Isoform 2 was not distinguished by RT-qPCR and was not analyzed on the protein level. Also, protein measurements performed by western blot are semi-quantitative and may not resemble the actual protein content in the cells. Nonetheless, despite these limitations, our study constitutes the most thorough overview of mRNA and protein regulation and spatial localisation of ACE2, and other SARS-CoV-2 related host molecules in differentiated primary human bronchial epithelial cells upon allergic and virus-induced inflammation in health and in asthma.

## Methods

### ALI culture

Primary human bronchial epithelial cells (Epithelix) were passaged in bronchial epithelial basal medium (BEBM, Lonza) with all supplements (Bronchial epithelial SingleQuots, Lonza). Cells were grown up to 80-90% confluency and then seeded into transwell inserts at a density of 1.5 x 10^5^ cells per well (0.40 μm pore size, 0.33 cm^2^, Corning). After reaching confluence, the apical medium was removed and the basal medium was changed to 1:1 BEBM (Lonza) with DMEM (Gibco), containing 0.06% wt/vol all-trans retinoic acid (Sigma). The medium was changed every 2-3 days and excess mucus was removed until cells were fully differentiated (^~^28days). HDM extract (Allergopharma) was applied apically on HBECs at a dose of 200 μg/ml of the protein concentration of the extract. Human IL-13 (Lubio Science) was diluted in OptiMem (ThermoFisher) and applied at 50ng/ml. Cells were kept at 37°C with 5% CO_2_.

### Viruses

Viral titer of human RV-A16 (Microbiologics Global Virology Center) was determined by plaque assay in H1-Hela cells (ATTC). Inactivation of virus was performed on UV-light at 254nm for 60min. Virus was diluted in OptiMem (ThermoFisher) and applied apically at a MOI 0.1 for two hours at 34,5°C with 5% CO_2_.

### RT-qPCR

RNA was isolated by RNeasy Plus Mini Kit (Qiagen). Yield and purity was measured by Nanodrop 2000 (ThermoFisher). RT was performed with RevertAid RT Reverse Transcription Kit (ThermoFisher). Quantitative PCR was performed with Maxima SYBR Green/ROX qPCR MasterMix (ThermoFisher) on Quantstudio 7 Real-Time PCR System (ThermoFisher). Relative quantification was calculated by 2^-ΔΔCt^ method described previously^49^. Sequences of primers used are summarized in the supplementary methods.

### Western blotting

Cells were lysed using RIPA buffer (ThermoFisher) containing cOmplete, Mini, EDTA-free protease inhibitor (Sigma). Protein concentration was determined by BCA Protein Assay kit (ThermoFisher). 10μg/sample with 4x Laemmli Buffer (Bio-Rad) containing β-mercaptoethanol was resolved in 20% Mini-PROTEAN TGX Gel (Witec AG). PNGase treatments have been performed according to manufacturer’s protocol (New England BioLabs). Gels were transblotted on nitrocellulose membrane (Advansta) using eBlot L1 (GenScript). Membranes were blocked and stained with primary antibody overnight in 5% nonfat dry milk in 0.1% PBST, visualized with HRP-conjugated secondary antibody using WesternBright Quantum (Advansta) and imaged by Fusion FX 7 Imaging System (Vilber). Antibodies used are rabbit anti-ACE2 (1:500, ab15348, Abcam), rabbit anti-ACE2 (1:1000, HPA000288, Sigma), rabbit anti-TMPRSS2 (1:1000, ab92323, Abcam), rabbit anti-NRP1 (1:50, HPA030278, Sigma), HRP-conjugated anti-rabbit (1:10000, 11-035-003, Jackson) and HRP anti β-actin (1:25000, ab49900, Abcam).

### IHC and Image Analysis

Cells were fixed in 4% PFA for 7min. Samples for cryosections have been embedded in *FSC22* section medium (Leica), cut at 6μm on a cryostat (Leica) and mounted on *SuperFrost Plus* glass slides (Menzel). Unreacted aldehydes were blocked by 0.1M glycine for 5 min. After two hours incubation in blocking medium containing 10% goat serum (Dako), 1% BSA (Sigma) and 0.2% TritonX-100 (Sigma), samples were stained overnight with primary antibody in diluted blocking solution (1:1 in PBS). Secondary antibodies were applied together with DAPI (Sigma) and Phalloidin-iFluor 633 (Abcam) for 2 hours at RT and then mounted with Fluoromount (Sigma). Primary Antibodies used were rabbit anti-ACE2 (1:500, ab15348, Abcam), rabbit anti-ACE2 (1:1000, HPA000288, Sigma), mouse anti-Occludin (1:200, OC-3F10, ThermoFisher), rabbit anti-TMPRSS2 (1:1000, ab92323, Abcam) and rabbit anti-NRP1 (1:50, HPA030278, Sigma). Rabbit Immunoglobin Fraction (X0936, Dako) and Mouse IgG1 Control (X0931, Dako) were diluted to same concentration as primary antibodies respectively as isotype controls. Goat anti-Mouse IgG Alexa 488 (1:1000, A11001, Invitrogen) and Goat anti-Rabbit IgG Alexa 546(1:500, A11010, Invitrogen) were used as secondary antibodies. Image acquisition was performed with LSM780 (Zeiss) by *ZEN* software and *ImageJ/Fiji* (NIH, Bethesda, USA) was used for image analysis.

### Transcriptome analyses

Next generation sequencing (NGS) from differentiated HBECs from control and patients with asthma, with or without IL-13 stimulation at 50ng/ml for 24 hours was performed as previously described^10^ and uploaded on the Gene Expression Omnibus (GEO) platform (https://www.ncbi.nlm.nih.gov/geo) and will be publicly available under the accession number GSE206510 upon publication. NGS from differentiated HBECs from control and patients with asthma, infected with RV-A16 were obtained from GEO platform under accession number GSE61161.

### Statistics

The data was analyzed by *GraphPad Prism* software (Version 9, San Diego, California, USA). P values of less than 0.05 (*), < 0.01 (**), < 0.001 (***) and < 0.0001 (****) were considered as significant. Statistics have been done comparing to vehicle, using Dunnett’s Multiple comparison on one way ANOVA if not stated differently.

## Author Contributions

Conceptualization: M.S., N.S., U.R. Methodology: N.S., U.R., M.D., M.H., P.W., G.T. Investigation, Visualization and Writing: N.S., M.S.

## Acknowledgements

This work was supported by the Swiss National Science Foundation (SNSF) grant (nr310030_189334/1) to MS.

