## Supplementary Information for "Regulation of mRNA transcripts, protein isoforms, glycosylation and spatial localization of ACE2 and other SARS-CoV-2-associated molecules in human airway epithelium upon viral infection and type 2 inflammation"

### Methods

#### Primary human bronchial epithelial cells (HBECs)

Primary human bronchial epithelial cells (HBECs) were isolated from control individuals and patients with asthma enrolled at the University Hospital, Jagiellonian University Medical College, Cracow, Poland, as described previously[1]. The study was granted ethical permission from Jagiellonian University Bioethics Committee, Poland (KBET/68/B/2008 and KBET/209/B/2011). All participants agreed to the study and provided a written, informed consent. Additionally, HBECs were purchased from two independent commercial sources: Lonza (Basel, Switzerland) and Epithelix (Plan-les-Ouates, Switzerland). The characteristics of HBECs used in the study are presented below in the Supplementary Table 1.

#### Air-liquid interface (ALI) cultures of bronchial epithelium

Primary human bronchial epithelial cells (HBECs) from passage 2 were cultured in the bronchial epithelial basal medium (BEBM) (CC-3171, Lonza, Basel, Switzerland) supplemented with the SingleQuot Kit (CC-4175, Lonza) in a humidified incubator at 37°C with 5% $CO_2$ for maximum 10 day, or until 80-90% confluency. Next, the cells were seeded on the transwell inserts with polyester membrane with 0.4 µm pore size (0.33 $cm^2$, Corning) at a density of $1.5x10^5$ cells per well in 24-well culture plates. After cells reached full confluence in the inserts (~2-5 days), apical medium was removed to enable differentiation of the cells in the air-liquid interface (ALI), as described previously[1]. Bronchial epithelial cells in ALI were differentiated in basolaterally supplemented BEBM medium (CC-3171, Lonza, Basel, Switzerland) containing all bullets except triiodothyronine (T3) and retinoic acid (RA) (CC-4175, Lonza) mixed 1:1 with DMEM (41965-039, Gibco) and fresh all-trans retinoic acid at 0.06% wt/vol (R2625, Sigma-Aldrich) in a humidified incubator at 37°C with 5% $CO_2$ for 21-28 days. Medium was changed every 2-3 days and excess mucus was removed. Experiments were performed after four weeks of differentiation in the ALI in OptiMem Medium (31985070, ThermoFisher).

**Reagents and viruses**

House dust mite extract (HDM) (Allergopharma, Reinbek, Germany) was diluted in the sterile 0.9% NaCl and stored in -20°C. The concentration of HDM used for the experiments was calculated according to the total protein content. HDM was applied apically at a dose of 200 μg/ml of protein. Human IL-13 (200-13, Lubio Science, Zürich, Switzerland) was diluted in OptiMem medium (31985070, ThermoFisher) and applied basolaterally on the HBECs at 50ng/ml. Human rhinovirus (RV-A16) was purchased from the Microbiologics Global Virology Center (Formerly Virapur). Inactivation of virus was performed by exposure of UV-light at 254nm for 60min at a distance of 2 cm. Virus was diluted in the OptiMem Medium (31985070, ThermoFisher) and applied apically on the HBECs at the multiplicity of infection (MOI) 0.1. Viral titer was determined by the standard plaque assay in H1-Hela cells bought from the ATCC. Briefly, H1-Hela cells were grown in six-well plates until reaching 80-90% confluency and subsequently infected with a 10-fold serial dilution of RV-A16 for 2 hours at 34.5°C with 5% $CO_2$, following an overlay medium containing 0.4% liquid agarose in DMEM. 2-3 days post-infection, the agar-overlay was removed, and cells were fixed with 4% PFA for 7 minutes. Cells were stained with 1% crystal violet solution (V5265, Sigma-Aldrich) and plaques were confirmed by inspection under a microscope.

**House dust mite, interleukin-13 stimulation, and rhinovirus infection model**

House dust mite (HDM) and interleukin-13 stimulations, followed by the human rhinovirus A16 (RV-A16) infection were performed in 200 μl apical and 600 μl basolateral Optimem medium (31985070, ThermoFisher). ALI-differentiated HBECs were treated apically with HDM (200 μg/ml of protein concentration) and basolaterally with IL-13 (50ng/ml), or vehicle, in the humidified incubator at 37°C with 5% $CO_2$. After 24 hours, cells were apically infected with RV-A16 at MOI 0.1 or treated with UV-RV-A16 at the same dose and cultured in a humidified incubator at 34.5°C with 5% $CO_2$. Next, 6 hours and 24 hours after infection cells were harvested and RNA and protein cellular lysates were collected and stored at -80°C. Some cells were fixed with 4% PFA (Fluka/Sigma Aldrich, Buch, Switzerland) for immunohistochemistry.

**Next-generation sequencing**

Next-generation sequencing (NGS) of the differentiated primary human bronchial epithelial cells from control subjects and patients with asthma, with or without IL-13 stimulation at 50ng/ml for 24 hours was performed as previously described[2]. Briefly, total RNA was extracted with an RNeasy Plus Micro Kit (Qiagen, Hilden, Germany), and samples with RNA integrity greater than 9.0 were chosen for library preparation with the TruSeq Stranded mRNA Sample Prep Kit (Ilumina, San Diego, California). Sequencing was performed on the Ilumina HiSeq 2500. The mRNA expression data are uploaded to the Gene Expression Omnibus platform (https://www.ncbi.nlm.nih.gov/geo) under the accession number GSE206510 and will be publicly available upon publication.

The mRNA expression data of HBECs from six control and six patients with asthma, infected in vitro with RV-A16 at the MOI10 for 24 hours have been previously published by Bai et al[3] and are publicly available at the Gene Expression Omnibus platform (https://www.ncbi.nlm.nih.gov/geo) under the accession number: GSE61141[3]. We reanalyzed these data regarding the expression of the SARS-CoV-2-related molecules.

**RT-qPCR**

RNA was isolated by RNaesy Plus Mini Kit (74034, Qiagen) according to the manufacturers protocol. Purity and yield were assessed by Nanodrop 2000 (ThermoFisher Scientific, Waltham, USA). Reverse transcription was performed using RevertAid RT Reverse Transcription Kit (K1691, ThermoFisher) according to the manufacturers recommendation. Quantstudio 7 Real-Time PCR System (ThermoFisher Scientific, Waltham, USA) was used for quantitative PCR, using Maxima SYBR Green/ROX qPCR Master Mix (K0221, ThermoFisher). Same amounts of cDNA per sample and condition were used. Gene expression of *long ACE2* (F: 5'-GGCGTAACGGACCCAGGAAAT-3', R: 5'-GGCCCATTGTCACCTTTGTGC-3'), *short ACE2* (F: 5'-GGCTACAAGTGCTTCATTGAGG-3', R: 5'-GGCCCATTGTCACCTTTGTGC-3') and RV-A16 positive strand (F: 5'-CGGGACTGCAAACACTACCT-3', R: 5'-CACCACGTGTGTCCCTAACA-3) was assessed in duplicates and normalized to elongation factor 1α (F: 5'-TGGTATTGGTACTGTTCCTG-3', R: 5'-CTTCACTCAAAGCTTCATGG-3'). Relative quantification was calculated by the $2^{-\Delta\Delta Ct}$ method described previously[4].

**Western blotting**

Cells were lysed with RIPA buffer (89901, ThermoFisher) containing cOmplete, Mini, EDTA-free protease inhibitor (46931590001, Sigma-Aldrich). Protein concentration of the cell lysates was determined by BCA Protein Assay kit (23225, ThermoFisher). PNGase (P0705S, New England BioLabs) treatments were performed under denaturing reaction conditions at 37°C for 1 hour according to the manufacturer's protocol. Same amounts of protein lysates (10µg) with 4x Laemmli sample buffer (1610747, Bio-Rad) containing β-mercaptoethanol (63689, Sigma-Aldrich) were resolved on 4-20% Mini-PROTEAN TGX Gel (M00656, Witec AG) using MOPS running buffer (M00138, Witec AG). Gels were transblotted on a nitrocellulose membrane (L-08006-100, Advansta) using eBlot L1 Wet Blotting Transfer System (GenScript, Leiden, Netherlands). The membranes were blocked with 5% nonfat dry milk in PBST (0.1% Tween-20 in 1x PBS) for 1 hour at room temperature (RT) and incubated with primary antibody diluted in blocking buffer over night at 4°C. Following three washing steps, membranes were incubated in horseradish peroxidase conjugated secondary antibody solution for 1 hour at RT. The membranes were then washed three times, developed using WesternBright Quantum (K-12042-D20, Advansta) and imaged by Fusion FX 7 Imaging System (Vilber, Collegien, France). Membranes were stripped with Restore PLUS Western Blot Stripping Buffer (46427, ThermoFisher) for 10 minutes and prior to subsequent staining with β-actin (1:25000, ab49900, Abcam). Band intensities were assessed by plotted areas using *ImageJ/Fiji* software. Intensity measurements have been first normalized to respective β-actin stained band and then normalized per percental intensity change of untreated control samples (Veh). Antibodies used are rabbit anti-ACE2 (1:500, ab15348, Abcam), rabbit anti-ACE2 (1:1000, HPA000288, Sigma), rabbit anti-TMPRSS2 (1:1000, ab92323, Abcam), rabbit anti-NRP1 (1:50, HPA030278, Sigma) and HRP-conjugated anti-rabbit (1:10000, 11-035-003, Jackson).

**Immunohistochemistry**

ALI-differentiated cells on inserts were fixed in 4% PFA in PBS for 7 min at RT and afterwards washed twice in 1 x PBS. The polyester membranes with cells were then removed from the transwell inserts. Samples for cryosections have been frozen in Clear Frozen Section Compound (FSC22, Leica), cut at a thickness of 6 µm in a cryostat (CM3050S, Leica) and mounted on the *SuperFrost Plus*™ glass slides (Menzel). Samples for the top-view acquisition have been processed free floating in 24-well culture dishes. Cryosections have been lined with PAP pen (NC9827128, ThermoFisher) and unreacted aldehydes were blocked by glycine (Sigma) at 0.1M in PBS for 5min. Samples have been incubated in blocking solution containing 10% normal goat serum (X0907, Dako), 1% bovine serum albumin (A3294, Sigma-Aldrich) and 0.2% TritonX-100 in PBS for 2 hours at RT. Primary antibodies have been diluted in blocking solution (1:1 in PBS) and incubated at 4°C overnight. Following three washing steps in 0.05% Tween20 in PBS, secondary antibodies together with DAPI in diluted blocking solution (1:1 in PBS) were applied for 2 hours at RT in the dark. Sections have been washed three times in 0.05%Tween20 in PBS before mounting with Fluoromount (F4680, Sigma Aldrich). Primary antibodies used were: rabbit anti-ACE2 (1:500, ab15348, Abcam), rabbit anti-ACE2 (1:1000, HPA000288, Sigma), mouse anti-Occludin (1:200, OC-3F10, ThermoFisher), rabbit anti-TMPRSS2 (1:1000, ab92323, Abcam) and rabbit anti-NRP1 (1:50, HPA030278, Sigma). Rabbit Immunoglobin Fraction (X0936, Dako) and mouse IgG1 Control (X0931, Dako) were diluted to the same concentration as primary antibodies respectively as isotype controls. Goat anti-Mouse IgG Alexa 488 (1:1000, A11001, Invitrogen) and Goat anti-Rabbit IgG Alexa 546(1:500, A11010, Invitrogen) were used as the secondary antibodies. Phallodin-iFluor 633 Reagent (1:1000, ab176758, Abcam) was used as structural marker and DAPI (1:1000, 10236276001, Sigma) to stain nuclei.

**Microscopy and Image Analysis**

Image acquisition was performed with Zeiss LSM780 and *ZEN* software, using a 40x objective. *ImageJ/Fiji* (NIH, Bethesda, USA) software was used for image analysis. Apical ACE2 (ab15348) intensity (Figure 3C) was measured on planar view on maximum intensity projection of apical Z-stacks (Figure 3D) up to and including tight junction layer, assessed by Occludin co-staining on 5 donors. On a randomly placed square on the same images, cell shape and area measurements have been performed by the human-assisted segmentation plugin *Cell Magic Wand Tool (*https://github.com/fitzlab/*CellMagicWand (February, 2022)).*

**Data analysis**

The data were analyzed by *GraphPad Prism* software (Version 9, San Diego, California, USA). P values of less than 0.05 (*), < 0.01 (**), < 0.001 (***) and < 0.0001 (****) were considered as significant. Statistics have been done comparing to the vehicle condition, using Dunnett´s Multiple comparison on One way ANOVA, if not stated differently.

Transcriptome data were processed as previously described[2]. Briefly, RNA-seq data were processed with the workflow available at https://github.com/uzh/ezRun. Significance threshold was set to fdr<0.05 calculated using the edgeR R package. N-linked glycosylation on aspargine (GO0006487) and SARS-CoV-2 associated molecules were curated from GSEA and MSigDB Database (Broad Institute, Massachusetts Institute of Technology, University of California, USA) and from literature (Figure S4). Analysis of biological processes and volcano plots were performed with use of Interactive ShinyApps available at https://fgcz-shiny.uzh.ch/.

**Supplementary Table 1. Clinical characteristics of the study subjects.**

| Gender | Pathology | Age Range | Origin | Cell source | iGCS (µg/d) | FEV1% predicted | Blood Eos (cells/1µl) |
|---|---|---|---|---|---|---|---|
| Female | Asthma | 51-60 | Caucasian | Epithelix | NA | NA | NA |
| Female | Asthma | 21-30 | Caucasian | Patients Cohort | 1000 | 77.9 | 395 |
| Female | Asthma | 51-60 | Caucasian | Patients Cohort | 500 | 109.3 | 1187 |
| Female | Asthma | 41-50 | Caucasian | Patients Cohort | 1000 | 112.3 | 590 |
| Female | Asthma | 31-40 | Caucasian | Patients Cohort | 1800 | 74.1 | 158 |
| Female | Asthma | 61-70 | Caucasian | Patients Cohort | 1000 | 94.9 | 259 |
| Female | Asthma | 51-60 | Caucasian | Patients Cohort | 1000 | 54.2 | 523 |
| Female | Asthma | 41-50 | Caucasian | Patients Cohort | 1000 | 74.8 | 356 |
| Female | Healthy | 10-20 | Caucasian | Epithelix | NA | NA | NA |
| Female | Healthy | 51-60 | "B" | Lonza | NA | NA | NA |
| Male | Healthy | 61-70 | Caucasian | Epithelix | NA | NA | NA |
| Male | Healthy | 51-60 | Caucasian | Epithelix | NA | NA | NA |
| Male | Healthy | 61-70 | Hispanic | Epithelix | NA | NA | NA |
| Male | Asthma | 51-60 | Caucasian | Epithelix | NA | NA | NA |
| Male | Asthma | 31-40 | Caucasian | Patients Cohort | 1000 | 98.5 | 90 |
| Male | Asthma | 51-50 | Caucasian | Patients Cohort | 500 | 87.7 | 848 |
| Male | Healthy | 71-80 | Caucasian | Epithelix | NA | NA | NA |
| Male | Healthy | 51-60 | Caucasian | Epithelix | NA | NA | NA |
| Male | Healthy | 71-80 | Caucasian | Epithelix | NA | NA | NA |
| Female | Healthy | NA | NA | Lonza | NA | NA | NA |

Primary human bronchial epithelial cells (HBECs) were obtained from the study subjects and commercial sources as specified. **iGCS**, inhaled glucocorticoids, recounted for flutikasone µg/d: **FEV1%**, forced expiratory volume: **Blood Eos**, eosinophils per 1µl of blood.

### Supplementary Figure 1

Signal Peptide · SARS-CoV-2 spike Interaction · Active site · **Unique sequence** · ADAM17 cleavage site · TMPRSS2 cleavage site · Transmembrane · *LC3-interacting region (LIR)* · SH2-binding(&Endocytic sorting signal · Phosphotyrosine-binding domain (PTB) · PDZ-binding

**Missmatch(---)** **Glycosylation** **Phosphorylation**

★ = Zn binding site    Peptidase_M2    Collectrin

```
               1                                                                             80
NP_001358344.1 MSSSSWLLLSLVAVTAAQSTIEEQAKTFLDKFNHEAEDLFYQSSLASWNYNTNITEENVQNMNNAGDKWSAFLKEQSTLA
NP_001373188.1 MSSSSWLLLSLVAVTAAQSTIEEQAKTFLDKFNHEAEDLFYQSSLASWNYNTNITEENVQNMNNAGDKWSAFLKEQSTLA
NP_001376331.1 MSSSSWLLLSLVAVTAAQSTIEEQAKTFLDKFNHEAEDLFYQSSLASWNYNTNITEENVQNMNNAGDKWSAFLKEQSTLA
NP_001375381.1 --------------------------------------------------------------------------------

               81                                                                           160
NP_001358344.1 QMYPLQEIQNLTVKLQLQALQQNGSSVLSEDKSKRLNTILNTMSTIYSTGKVCNPDNPQECLLLEPGLNEIMANSLDYNE
NP_001373188.1 QMYPLQEIQNLTVKLQLQALQQNGSSVLSEDKSKRLNTILNTMSTIYSTGKVCNPDNPQECLLLEPGLNEIMANSLDYNE
NP_001376331.1 QMYPLQEIQNLTVKLQLQALQQNGSSVLSEDKSKRLNTILNTMSTIYSTGKVCNPDNPQECLLLEPGLNEIMANSLDYNE
NP_001375381.1 --------------------------------------------------------------------------------

               161                                                                          240
NP_001358344.1 RLWAWESWRSEVGKQLRPLYEEYVVLKNEMARANHYEDYGDYWRGDYEVNGVDGYDYSRGQLIEDVEHTFEEIKPLYEHL
NP_001373188.1 RLWAWESWRSEVGKQLRPLYEEYVVLKNEMARANHYEDYGDYWRGDYEVNGVDGYDYSRGQLIEDVEHTFEEIKPLYEHL
NP_001376331.1 RLWAWESWRSEVGKQLRPLYEEYVVLKNEMARANHYEDYGDYWRGDYEVNGVDGYDYSRGQLIEDVEHTFEEIKPLYEHL
NP_001375381.1 --------------------------------------------------------------------------------

               241                                                                          320
NP_001358344.1 HAYVRAKLMNAYPSYISPIGCLPAHLLGDMWGRFWTNLYSLTVPFGQKPNIDVTDAMVDQAWDAQRIFKEAEKFFVSVGL
NP_001373188.1 HAYVRAKLMNAYPSYISPIGCLPAHLLGDMWGRFWTNLYSLTVPFGQKPNIDVTDAMVDQAWDAQRIFKEAEKFFVSVGL
NP_001376331.1 HAYVRAKLMNAYPSYISPIGCLPAHLLGDMWGRFWTNLYSLTVPFGQKPNIDVTDAMVDQAWDAQRIFKEAEKFFVSVGL
NP_001375381.1 --------------------------------------------------------------------------------

               321                                                   ★   ★              400
NP_001358344.1 PNMTQGFWENSMLTDPGNVQKAVCHPTAWDLGKGDFRILMCTKVTMDDFLTAHHEMGHIQYDMAYAAQPFLLRNGANEGF
NP_001373188.1 PNMTQGFWENSMLTDPGNVQKAVCHPTAWDLGKGDFRILMCTKVTMDDFLTAHHEMGHIQYDMAYAAQPFLLRNGANEGF
NP_001376331.1 PNMTQGFWENSMLTDPGNVQKAVCHPTAWDLGKGDFRILMCTKVTMDDFLTAHHEMGHIQYDMAYAAQPFLLRNGANEGF
NP_001375381.1 --------------------------MREAGWDKGGRILMCTKVTMDDFLTAHHEMGHIQYDMAYAAQPFLLRNGANEGF

            401 ★                                                                           480
NP_001358344.1 HEAVGEIMSLSAATPKHLKSIGLLSPDFQEDNETEINFLLKQALTIVGTLPFTYMLEKWRWMVFKGEIPKDQWMKKWWEM
NP_001373188.1 HEAVGEIMSLSAATPKHLKSIGLLSPDFQEDNETEINFLLKQALTIVGTLPFTYMLEKWRWMVFKGEIPKDQWMKKWWEM
NP_001376331.1 HEAVGEIMSLSAATPKHLKSIGLLSPDFQEDNETEINFLLKQALTIVGTLPFTYMLEKWRWMVFKGEIPKDQWMKKWWEM
NP_001375381.1 HEAVGEIMSLSAATPKHLKSIGLLSPDFQEDNETEINFLLKQALTIVGTLPFTYMLEKWRWMVFKGEIPKDQWMKKWWEM

               481                                                                          560
NP_001358344.1 KREIVGVVEPVPHDETYCDPASLFHVSNDYSFIRYYTRTLYQFQFQEALCQAAKHEGPLHKCDISNSTEAGQKLFNMLRL
NP_001373188.1 KREIVGVVEPVPHDETYCDPASLFHVSNDYSFIRYYTRTLYQFQFQEALCQAAKHEGPLHKCDISNSTEAGQKLFNMLRL
NP_001376331.1 KREIVGVVEPVPHDETYCDPASLFHVSNDYSFIRYYTRTLYQFQFQEALCQAAKHEGPLHKCDISNSTEAGQKLL-----
NP_001375381.1 KREIVGVVEPVPHDETYCDPASLFHVSNDYSFIRYYTRTLYQFQFQEALCQAAKHEGPLHKCDISNSTEAGQKLFNMLRL

               561                                                                          640
NP_001358344.1 GKSEPWTLALENVVGAKNMNVRPLLNYFEPLFTWLKDQNKNSFVGWSTDWSPYADQSIKVRISLKSALGDKAYEWNDNEM
NP_001373188.1 GKSEPWTLALENVVGAKNMNVRPLLNYFEPLFTWLKDQNKNSFVGWSTDWSPYADQSIKVRISLKSALGDKAYEWNDNEM
NP_001376331.1 --------------------------------------------------------------------------------
NP_001375381.1 GKSEPWTLALENVVGAKNMNVRPLLNYFEPLFTWLKDQNKNSFVGWSTDWSPYADQSIKVRISLKSALGDKAYEWNDNEM

               641                                                                          720
NP_001358344.1 YLFRSSVAYAMRQYFLKVKNQMILFGEEDVRVANLKPRISFNFFVTAPKNVSDIIPRTEVEKAIRMSRSRINDAFRLNDN
NP_001373188.1 YLFRSSVAYAMRQYFLKVKNQMILFGEEDVRVANLKPRISFNFFVTAPKNVSDIIPRTEVEKAIRMSRSRINDAFRLNDN
NP_001376331.1 --------------------------EEDVRVANLKPRISFNFFVTAPKNVSDIIPRTEVEKAIRMSRSRINDAFRLNDN
NP_001375381.1 YLFRSSVAYAMRQYFLKVKNQMILFGEEDVRVANLKPRISFNFFVTAPKNVSDIIPRTEVEKAIRMSRSRINDAFRLNDN

               721                                                                          800
NP_001358344.1 SLEFLGIQPTLGPPNQPPVSIWLIVFGVVMGVIVVGIVILIFTGIRDRKKKNKARSGENPYASIDISKGENNPGFQNTDD
NP_001373188.1 SLEFLGIQPTLGPPNQPPVSIWLIVFGVVMGVIVVGIVILIFTGIRDRKKPTPLLGKSWLTAILKD--------------
NP_001376331.1 SLEFLGIQPTLGPPNQPPVSIWLIVFGVVMGVIVVGIVILIFTGIRDRKKKNKARSGENPYASIDISKGENNPGFQNTDD
NP_001375381.1 SLEFLGIQPTLGPPNQPPVSIWLIVFGVVMGVIVVGIVILIFTGIRDRKKKNKARSGENPYASIDISKGENNPGFQNTDD

               801
NP_001358344.1 VQTSF
NP_001373188.1 -----
NP_001376331.1 VQTSF
NP_001375381.1 VQTSF
```

The amino acid sequences of NP_001358344.1 and NP_068576.1 are equal, therefore only NP_001358344.1 is shown.

The amino acid sequences of NP_001376331.1 and NP_001373189.1 are equal, therefore only NP_001376331.1 is shown.

**Supplementary Figure 1**

Full sequence alignments of ACE2 isoform precursors, showing functional and structural domains in the protein sequence. Angiotensin-converting enzyme 2 Isoform 1 precursor is encoded by the transcript variants 1 (NP_001358344.1) and 2 (NP_068576.1). Isoform 2 precursor is encoded by the transcript variant 3 (NP_001373188.1). Isoform 3 precursor is encoded by the transcript variants 4 (NP_001373189.1) and 6 (NP_00137633.1). Isoform 4 precursor is encoded by the transcript variant 5 (NP_001375381.1). The amino-acid sequences of NP_001358344.1 and NP_068576.1 are equal and sequences of NP_001376331.1 and NP_001375381.1 are equal, therefore, NP_068576.1 and NP_001375381.1 are not shown. Source of sequences: NCBI[5].

Supplementary Figure 2

**a**

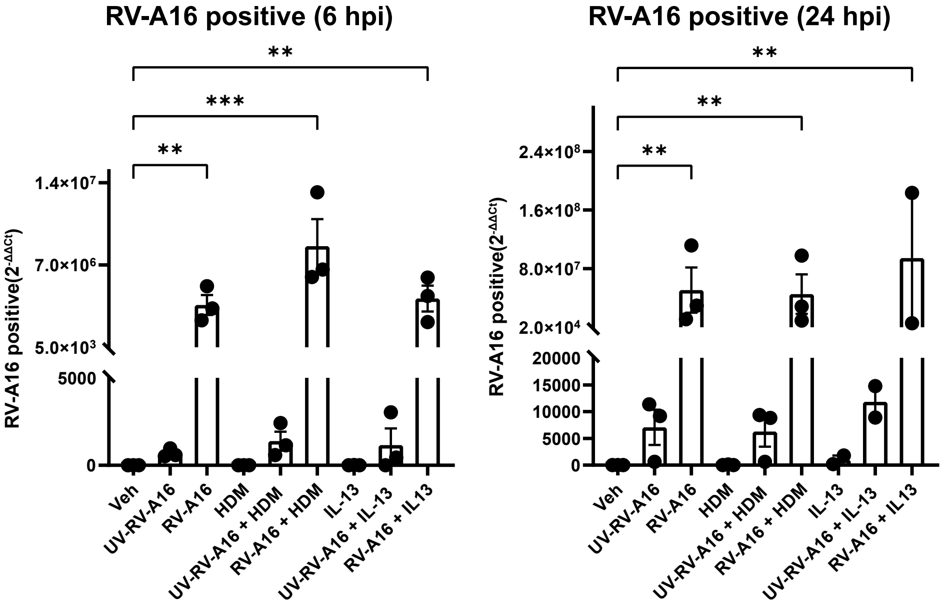

**Supplementary Figure 2**

Relative expression of RV-A16 at 6 and 24 hours post-infection (hpi). RV-A16 positive-strand were assessed in duplicates and normalized to elongation factor 1α, shown with $2^{-\Delta\Delta CT}$ values to unstimulated condition (Veh). Individual values from each subject are shown by dots; bars represent mean ± SEM. Statistics was performed using Kruskal-Wallis (ANOVA). **HDM**, house dust mite; **hpi**, hours post-infection; **Veh**, vehicle; **RV-A16**, human rhinovirus A16; **UV-RV-A16**, UV-light inactivated human rhinovirus A16.

Supplementary Figure 3

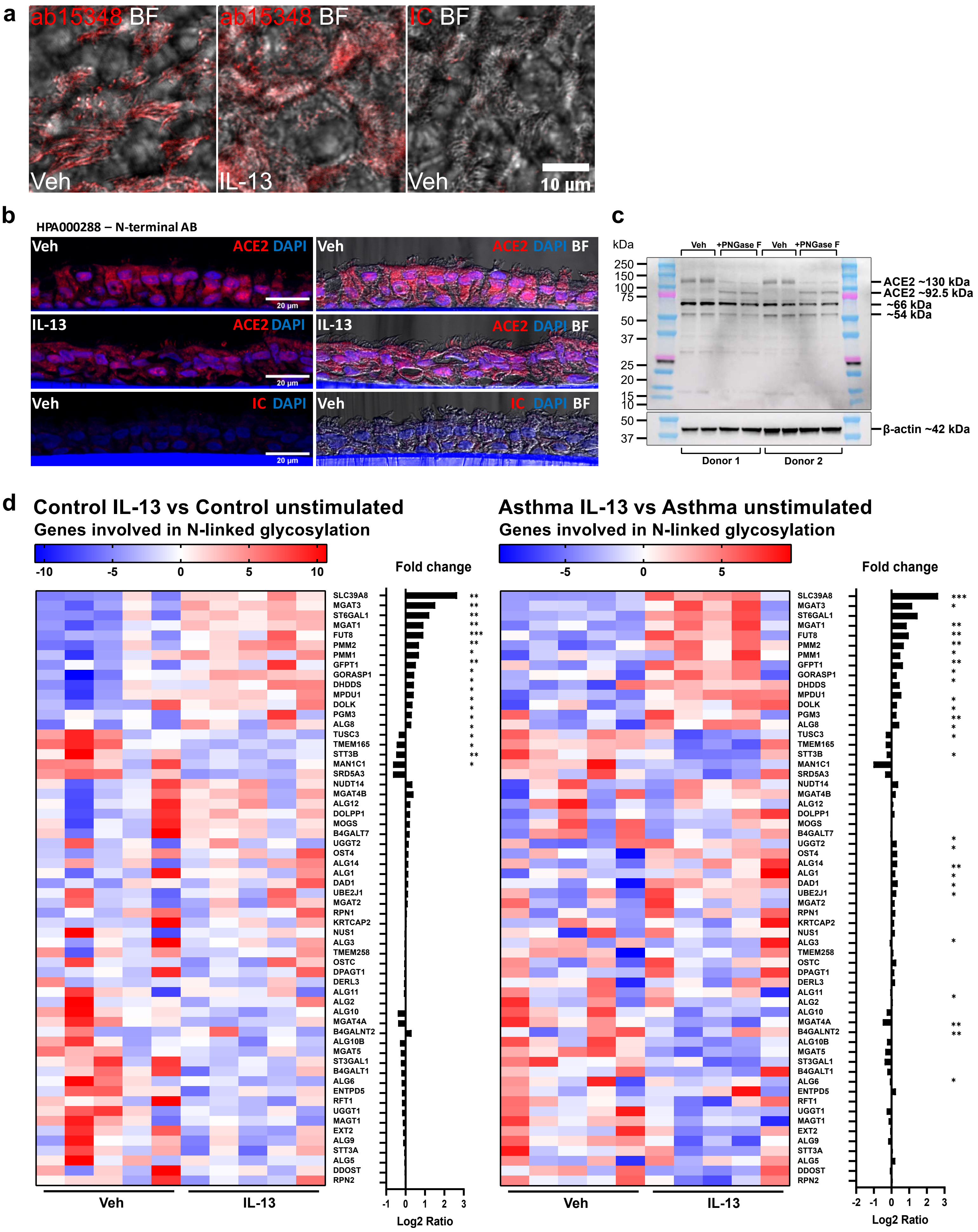

**Supplementary Figure 3**

**a)** Representative confocal microscopy of apical planar view of C-terminal ACE2 antibody (ab15348) staining of bronchial epithelial cilia in vehicle and IL-13 treated samples. **b)** Representative confocal microscopy of transversal view on cryosections from vehicle and IL-13 treated primary HBECs, stained with N-terminal ACE2 antibody (HPA000288). **c)** Representative western blot image of ACE2 expression detected in primary HBECs with anti-ACE2 (HPA000288) on two donors with PNGase treated lysates in duplicates. **d)** Heatmaps and fold changes of expression of all genes in N-linked glycosylation pathway (GO: 0006487) upon IL-13 treatment in control (n=5), left, and asthma (n=5), right. **IC**, isotype control; **BF**, bright field; **Veh**, vehicle.

Supplementary Figure 4

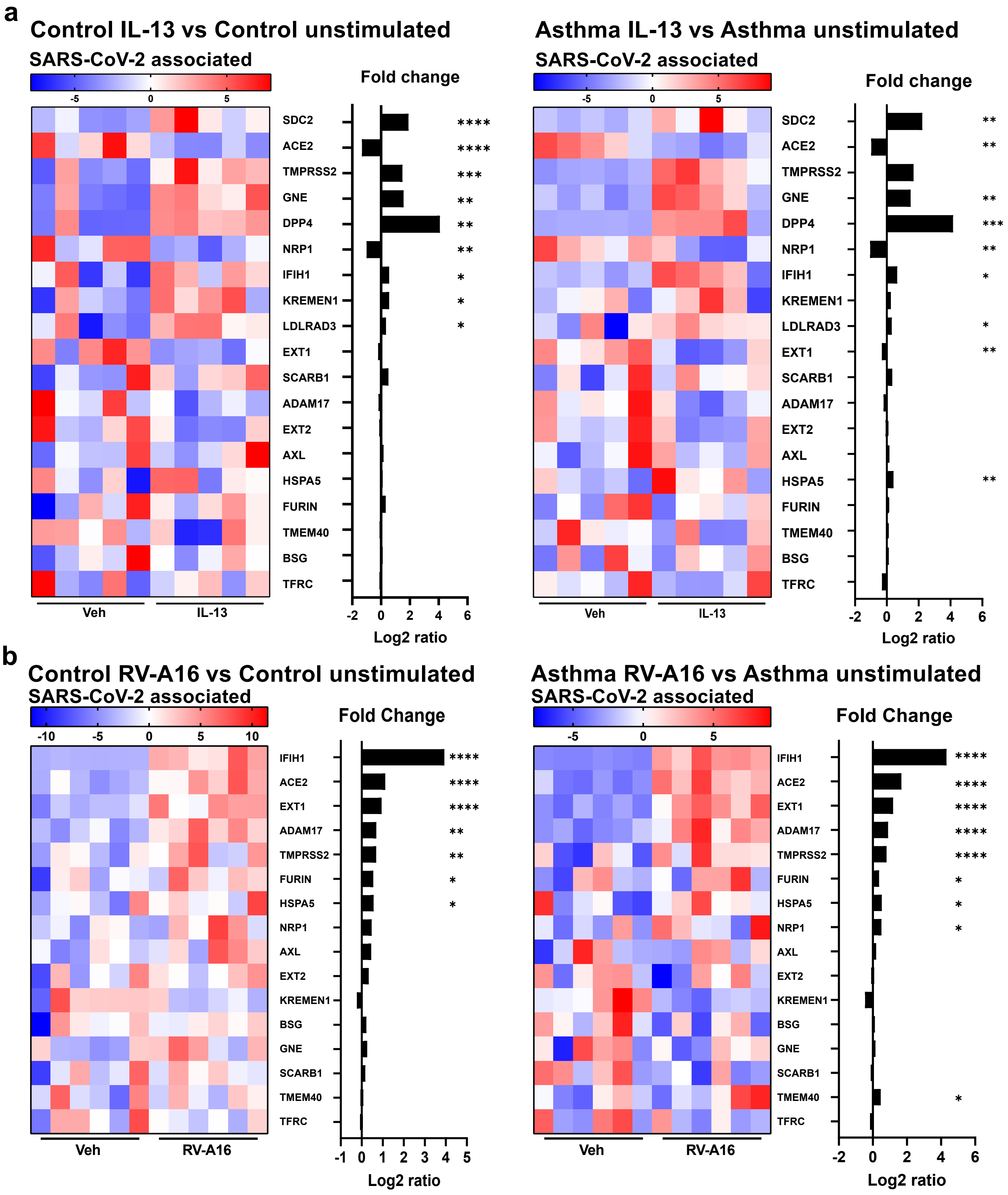


c

| GENE | Comments |
| --- | --- |
| ACE2, NRP1, SCARB1, AXL, BSG, LDLRAD3, TMEM40, CLEC4G, CD209, CLEC4M, ASGR1, KREMEN1, TFRC, HSPA5 | Candidate Receptors, as reviewed in Baggen et al. (2021) |
| SDC2, EXT1, EXT2 | Heparan Sulfate biosynthesis (Kreuger et al. (2012), Hudak et al. (2021) and Liu et al. (2021)) |
| GNE | Sialic acid precursor (Chen et al. (2016)) |
| TMPRSS2, ADAM17, FURIN | Proteases (Radzikowksa et al. (2020)) |
| DPP4 | Zhang et al. (2021) |
| IFIH1 | Yin et al. (2021) |

**Supplementary Figure 4**

**a)** Heatmap and fold change (log2FC expression) of all investigated SARS-CoV-2 associated genes upon IL-13 treatment in HBEC from controls (n=5) and asthma (n=5). Not detected genes are not shown. **b)** Heatmaps and fold changes (log2FC expression) of all investigated SARS-CoV-2 associated genes upon RV-A16 infection in HBEC from control (n=6) and asthma (n=6). **c)** Table summarizing SARS-CoV-2 associated genes and references. **RV-A16**, human rhinovirus A16.
